## supplementary tables and figures for "Hierarchical Patterns of Soil Nematode Biodiversity in the Atacama Desert: Insights Across Biological Scales"

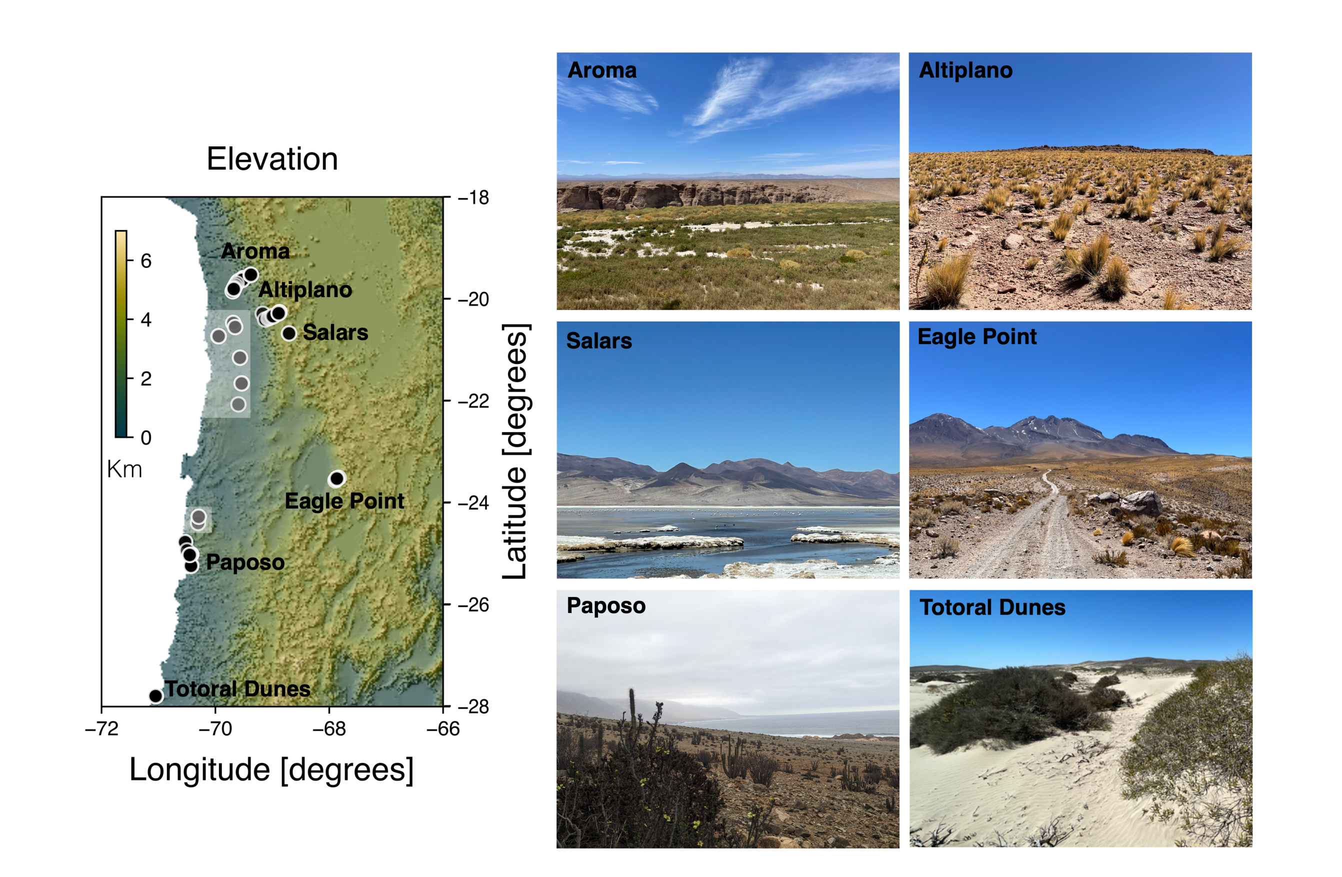


**Fig. S1.** Transects and sampling spots distributed across the Atacama Desert, shading corresponds to elevation in Km above sea level. Sampling spots in white shadow boxes represent no formal transects, and areas where no nematodes were found. Pictures for each of the transects highlight the differences in the ecosystems. The Aroma transect is located near the Quebrada de Aroma, a fresh water stream and vegetation coverage is high. The Altiplano transect is a high altitude transect with lower vegetation and water streams with seasonal rainfall. The Salars transect(including Salar de Huasco and Laguna Grande) correspond to wetlands with high salinity patches and fresh water inflow. The Eagle Point transect is a high altitude transect with lower vegetation. The Paposo transect is close the coast and receives moisture in the form of fog and has high plant endemism. The Totoral Dunes transect is also located close to the cost and is characterized by sandy dunes.

**Table S1. Sampling coordinates of Totoral Dunes (TDT) locality**

| Locality | Longitud | Latitude |
|---|---|---|
| TDT | -71,04472222 | -27,79444444 |
| TDT | -71,04472222 | -27,79416667 |
| TDT | -71,045 | -27,79416667 |
| TDT | -71,04527778 | -27,79416667 |
| TDT | -71,04555556 | -27,79416667 |
| TDT | -71,04611111 | -27,79416667 |
| TDT | -71,04638889 | -27,79416667 |
| TDT | -71,04694444 | -27,79388889 |
| TDT | -71,04722222 | -27,79388889 |
| TDT | -71,04777778 | -27,79388889 |
| TDT | -71,04805556 | -27,79388889 |
| TDT | -71,04833333 | -27,79388889 |
| TDT | -71,04888889 | -27,79361111 |
| TDT | -71,04916667 | -27,79361111 |
| TDT | -71,04944444 | -27,79361111 |
| TDT | -71,04944444 | -27,79361111 |
| TDT | -71,04972222 | -27,79333333 |
| TDT | -71,05027778 | -27,79361111 |
| TDT | -71,05027778 | -27,79333333 |
| TDT | -71,05055556 | -27,79333333 |

**Table S2. Sampling coordinates of Paposo (PAP) locality**

| Locality | Longitude | Latitude |
|---|---|---|
| PAP | -70,423056 | -25,014167 |
| PAP | -70,423056 | -25,014167 |
| PAP | -70,425833 | -25,011667 |
| PAP | -70,428333 | -25,012778 |
| PAP | -70,429444 | -25,018333 |
| PAP | -70,532222 | -24,772778 |
| PAP | -70,530833 | -24,7725 |
| PAP | -70,498056 | -24,935 |
| PAP | -70,425 | -25,233889 |
| PAP | -70,425278 | -25,235 |
| PAP | -70,426389 | -25,235556 |
| PAP | -70,427778 | -25,236944 |
| PAP | -70,445 | -25,021389 |
| PAP | -70,450833 | -25,024444 |
| PAP | -70,451111 | -25,025 |

**Table S3. Sampling coordinates of Eagle Point transect (EPT) locality**

| Locality | Longitude | Latitude |
|---|---|---|
| EPT | -67,892222 | -23,563056 |
| EPT | -67,877222 | -23,541944 |
| EPT | -67,860556 | -23,521667 |
| EPT | -67,848889 | -23,516111 |
| EPT | -67,834444 | -23,510556 |
| EPT | -67,865278 | -23,526667 |
| EPT | -67,865278 | -23,526667 |
| EPT | -67,865278 | -23,526667 |
| EPT | -67,865278 | -23,526667 |
| EPT | -67,865278 | -23,526667 |
| EPT | -67,865278 | -23,526667 |
| EPT | -67,865278 | -23,526667 |
| EPT | -67,865278 | -23,526667 |
| EPT | -67,865278 | -23,526667 |
| EPT | -67,865278 | -23,526667 |
| EPT | -67,865278 | -23,526667 |
| EPT | -67,865278 | -23,526667 |
| EPT | -67,865278 | -23,526667 |
| EPT | -67,865278 | -23,526667 |
| EPT | -67,865278 | -23,526667 |
| EPT | -67,865278 | -23,526667 |

**Table S4. Sampling coordinates of Aroma (ARO) locality**

| Locality | Longitude | Latitude |
|---|---|---|
| ARO | -69,677778 | -19,807222 |
| ARO | -69,677778 | -19,807222 |
| ARO | -69,677778 | -19,807222 |
| ARO | -69,608611 | -19,665833 |
| ARO | -69,591944 | -19,654722 |
| ARO | -69,581667 | -19,644722 |
| ARO | -69,507222 | -19,595833 |
| ARO | -69,510556 | -19,595278 |
| ARO | -69,510556 | -19,595278 |
| ARO | -69,381111 | -19,528611 |
| ARO | -69,381111 | -19,528611 |
| ARO | -69,3775 | -19,53 |
| ARO | -69,378611 | -19,528056 |
| ARO | -69,374167 | -19,526667 |
| ARO | -69,526111 | -19,625556 |
| ARO | -69,526111 | -19,625556 |
| ARO | -69,528889 | -19,626389 |
| ARO | -69,626111 | -19,697778 |
| ARO | -69,648889 | -19,739167 |
| ARO | -69,667222 | -19,781111 |
| ARO | -69,695 | -19,849722 |
| ARO | -69,695 | -19,849722 |
| ARO | -69,696389 | -19,849722 |

**Table S5. Sampling coordinates of Altiplano (ALT) locality**

| Locality | Longitude | Latitude |
|---|---|---|
| ALT | -69,126667 | -20,405278 |
| ALT | -69,069722 | -20,381389 |
| ALT | -69,042778 | -20,369167 |
| ALT | -69,010556 | -20,355 |
| ALT | -68,975278 | -20,339722 |
| ALT | -68,991944 | -20,339722 |
| ALT | -69,168333 | -20,291667 |

**Table S6. Sampling coordinates of the Salars locality**

| Locality | Longitude | Latitude |
|---|---|---|
| Salars | -68,875556 | -20,2625 |
| Salars | -68,875556 | -20,2625 |
| Salars | -68,886984 | -20,276002 |
| Salars | -68,886959 | -20,275679 |
| Salars | -68,875961 | -20,26247 |
| Salars | -68,875839 | -20,262499 |
| Salars | -68,889651 | -20,282766 |
| Salars | -68,889289 | -20,28312 |
| Salars | -68,889919 | -20,282545 |
| Salars | -68,706111 | -20,678333 |
| Salars | -68,705833 | -20,678611 |

**Table S7. Sampling coordinates in areas outside the defined sampling localities.**

| Locality | Longitude | Latitude |
|---|---|---|
| TAM | -69,686389 | -20,472778 |
| TAM | -69,686389 | -20,472778 |
| TAM | -69,653333 | -20,553611 |
| TAM | -69,653611 | -20,553056 |
| RL | -69,535556 | -21,658056 |
| RL | -69,540278 | -21,66 |
| MET | -69,595833 | -22,070556 |
| CDM | -70,304444 | -24,398611 |
| CDM | -70,293333 | -24,275556 |
| CDM | -70,293333 | -24,275556 |
| BTM | -69,568333 | -21,154167 |
| BTM | -69,568333 | -21,154167 |
| acCPAJ | -69,945556 | -20,732778 |
| acCPAJ | -69,945278 | -20,7325 |
| acCPAJ | -69,941667 | -20,7325 |

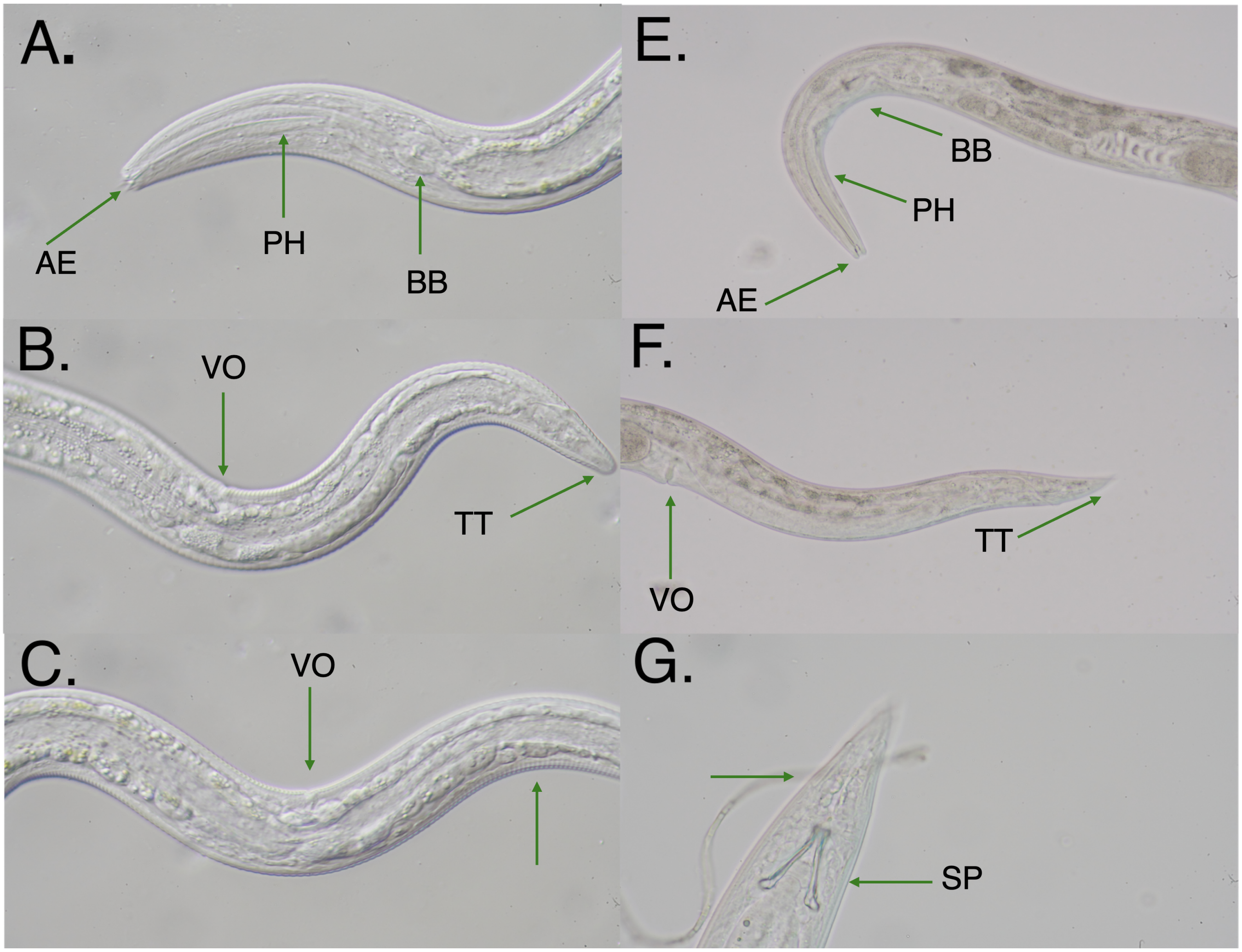


**Fig. S2.** Morphological characters used for the identification of extracted nematodes to families and genera, as exemplified by *Acrobeloides* (A-C) and *Panagrolaimus* (D-F). In panel A, a complex labial region can be seen (AE), and in E a simple labial region . *Diagnostic features: shape and size of stoma, presence or absence of tooth or odontostyle and labial structure.* In A and E both cases the presence of a well developed muscular Pharynx (PH) followed by a basal bulb with a grinder (BB). *Diagnostic features: musculature, shape of the pharynx and presence or absence and structure of bulb.* In B and F the vulva opening is indicated (VO) in female individuals in both cases, the tail tip differs (TT) between B and F, round and conoid tail, respectively. *Diagnostic features: reproductive structures and its position relative to whole body length, number of gonads, shape and length of tail.* In G a male individual is seen identified by the presence of a spicule (SP), in contrast the identified species in A-C only displayed females individuals as seen from morphological structures (VO) and experimental work, additionally in C a coarsely annulated cuticle is seen and in G a finely annulated cuticle. *Diagnostic features: presence and shape of spicule and cuticle structure*

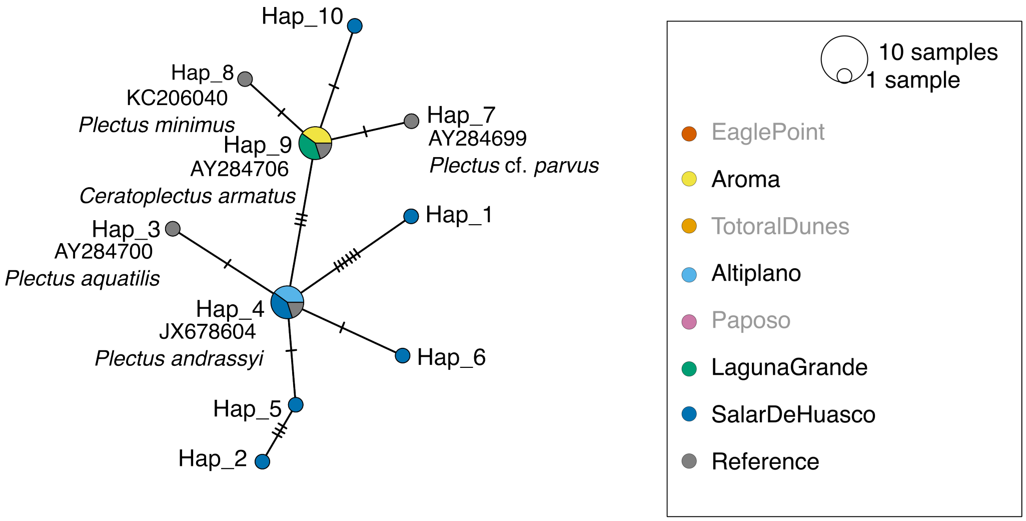


**Fig. S3.** Haplotype network for Plectidae. Based on 18S rRNA sequences. Color refers to sampling locations. Locations in grey are not present. Reference sequences were retrieved from the curated 18S-NemaBase (1). Accession number and species name is written next to the respective node of haplotype (abbreviated to "Hap"). Vertical hatchmarks indicate the number of differences from one node to the next.

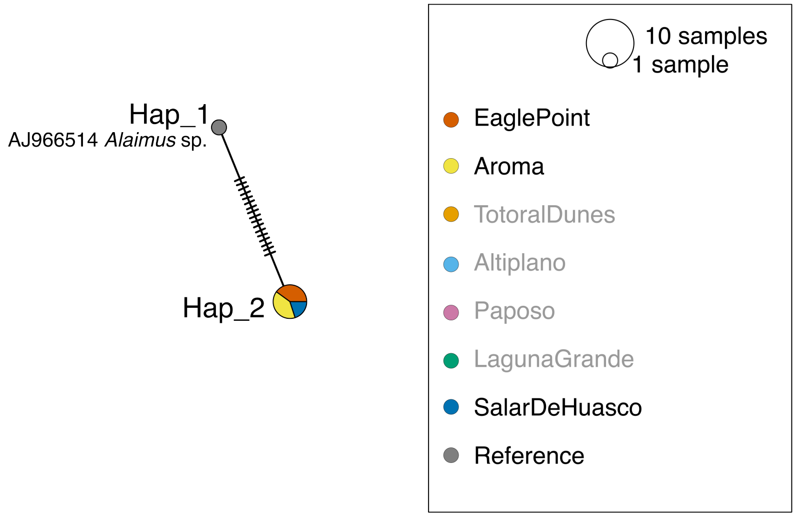


**Fig. S4.** Haplotype network for Alaimidae. Based on 18S rRNA sequences. Color refers to sampling locations. Locations in grey are not present. Reference sequences were retrieved from the curated 18S-NemaBase (1). Accession number and species name is written next to the respective node of haplotype (abbreviated to "Hap"). Vertical hatchmarks indicate the number of differences from one node to the next.

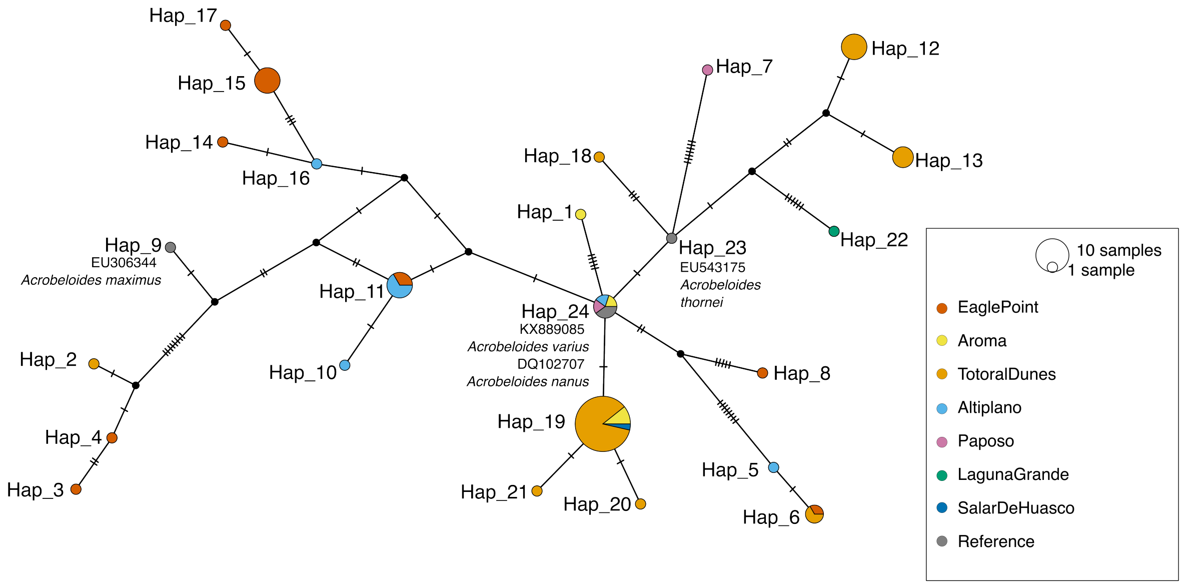


**Fig. S5.** Haplotype network for *Acrobeloides*. Based on 18S rRNA sequences. Color refers to sampling locations. Reference sequences were retrieved from the curated 18S-NemaBase (1). Accession number and species name is written next to the respective node of haplotype (abbreviated to "Hap"). Vertical hatchmarks indicate the number of differences from one node to the next.

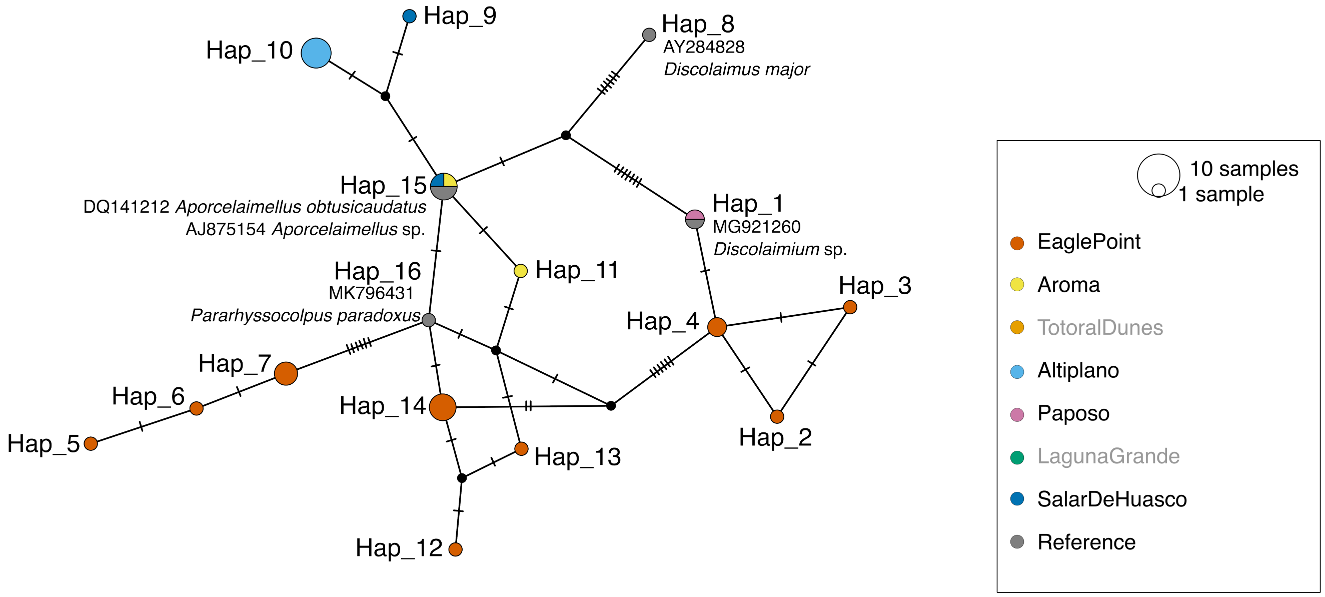


**Fig. S6.** Haplotype network for Dorylamida. Based on 18S rRNA sequences. Color refers to sampling locations. Locations in grey are not present. Reference sequences were retrieved from the curated 18S-NemaBase (1). Accession number and species name is written next to the respective node of haplotype (abbreviated to "Hap"). Vertical hatchmarks indicate the number of differences from one node to the next.

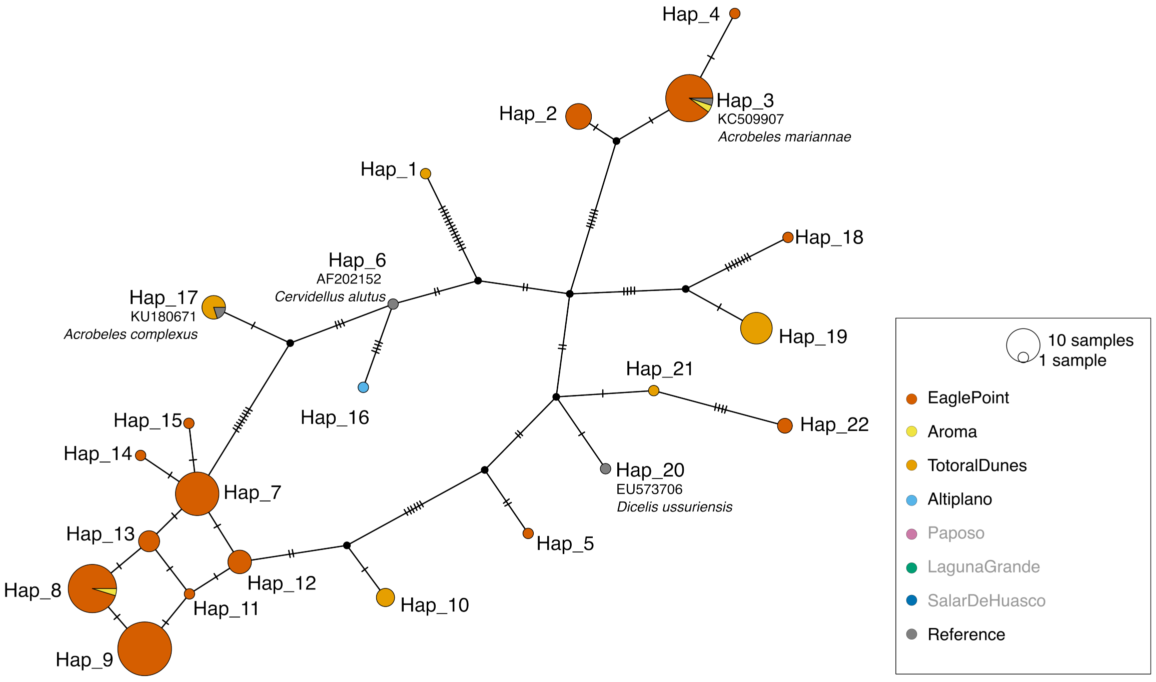


**Fig. S7.** Haplotype network for *Acrobeles*. Based on 18S rRNA sequences. Color refers to sampling locations. Locations in grey are not present. Reference sequences were retrieved from the curated 18S-NemaBase (1). Accession number and species name is written next to the respective node of haplotype (abbreviated to "Hap"). Vertical hatchmarks indicate the number of differences from one node to the next.

**Table S8. Details on the curated alignments used for haplotype networks, total nucleotide diversity ($\pi$) and total genetic diversity (Watterson's estimator, $\theta$).**

| Taxonomic group | Number of sequences | Alignment length (bp) | Number of segregating sites |
|---|---|---|---|
| Plectidae | 18 | 533 | 18 |
| Alaimidae | 6 | 497 | 17 |
| *Acrobeloides* | 75 | 487 | 51 |
| Dorylamida | 30 | 421 | 24 |
| *Acrobeles* | 129 | 328 | 47 |
| *Panagrolaimus* | 47 | 367 | 67 |
| Aphelenchoidea | 20 | 645 | 166 |

**Table S9. Details on nucleotide diversity ($\pi$) and genetic diversity (Watterson's estimator, $\theta$) for each taxonomic group based on every location. The number of sequences and number of haplotypes per location are also specified. The geographic distance of the different regions was tested against the genetic distance (Euclidean distance) with Mantel statistics based on Spearman's rank correlation ($\rho$). The references were excluded from the statiscal analyses (grey column). Significant p-values (p <0.05) are marked with an asterix.**

| Taxonomic group | Measurement | Eagle Point | Aroma | Totoral Dunes | Altiplano | Paposo | Salars | Reference | Mantel statistics |
|---|---|---|---|---|---|---|---|---|---|
| Plectidae | $\pi$ | NA | 0 | NA | 0 | NA | 0.0082 | 0.00568 | r = 0,<br>p = 0.66667,<br>5 permutations |
| | $\theta$ | NA | 0 | NA | 0 | NA | 0.0104 | 0.00545 | |
| | Haplotypes | NA | 1 | NA | 1 | NA | 7 | 5 | |
| | Sequences | 0 | 2 | 0 | 2 | 0 | 9 | 5 | |
| Alaimidae | $\pi$ | 0 | 0 | NA | NA | NA | NA | NA | NA |
| | $\theta$ | 0 | 0 | NA | NA | NA | NA | NA | |
| | Haplotypes | 1 | 1 | NA | NA | NA | 1 | 1 | |
| | Sequences | 2 | 2 | 0 | 0 | 0 | 1 | 1 | |
| *Acrobeloides* | $\pi$ | 0.01938 | 0.0063 | 0.00932 | 0.00959 | 0.0186 | 0.0186 | 0.00723 | r = -0.07527,<br>p = 0.55278,<br>719 permutations |
| | $\theta$ | 0.01935 | 0.007 | 0.01457 | 0.01195 | 0.0186 | 0.0186 | 0.00789 | |
| | Haplotypes | 8 | 3 | 8 | 5 | 2 | 2 | 3 | |
| | Sequences | 14 | 5 | 40 | 8 | 2 | 2 | 4 | |
| Dorylamida | $\pi$ | 0.01788 | 0.0024 | NA | 0 | NA | 0.0048 | 0.01449 | r = 0.7143,<br>p = 0.16667,<br>23 permutations |
| | $\theta$ | 0.0112 | 0.0024 | NA | 0 | NA | 0.0048 | 0.01596 | |
| | Haplotypes | 9 | 2 | NA | 1 | 1 | 2 | 4 | |
| | Sequences | 15 | 2 | 0 | 5 | 1 | 2 | 5 | |
| *Acrobeles* | $\pi$ | 0.02342 | 0.0491 | 0.03136 | NA | NA | NA | 0.0317 | r = -0.5,<br>p = 0.83333,<br>5 permutations |
| | $\theta$ | 0.0194 | 0.0491 | 0.02943 | NA | NA | NA | 0.03012 | |
| | Haplotypes | 14 | 2 | 5 | 1 | NA | NA | 4 | |
| | Sequences | 104 | 2 | 18 | 1 | 0 | 0 | 4 | |
| *Panagrolaimus* | $\pi$ | 0.06548 | 0.055 | 0 | 0.05476 | 0.03217 | NA | 0.04137 | r = 0.6,<br>p = 0.041667 *,<br>119 permutations |
| | $\theta$ | 0.06144 | 0.055 | 0 | 0.04056 | 0.02913 | NA | 0.04603 | |
| | Haplotypes | 3 | 3 | 1 | 8 | 5 | NA | 4 | |
| | Sequences | 4 | 3 | 5 | 15 | 13 | 0 | 5 | |
| Aphelenchoidea | $\pi$ | 0.05241 | 0.0634 | NA | NA | NA | NA | 0.1135 | NA |
| | $\theta$ | 0.04538 | 0.0836 | NA | NA | NA | NA | 0.09555 | |
| | Haplotypes | 5 | 5 | NA | 1 | 1 | NA | 6 | |
| | Sequences | 6 | 6 | 0 | 1 | 1 | 0 | 6 | |

**Table S10. Summary of barcoding data (18S rRNA). Sorted by locality and assigned group for TCS haplotype networks. Number of haplotype per group and locality is given, supported by the assigned haplotype number of the networks (Hap). Molecular based identification according to best hit (lowest E-value and a high percentage of identity) against the curated 18S-NemaBase is specified.**

| Locality | Haplotype ID | Count Haplotypes | Haplotypes (Hap) | Molecular Identification | Count Families | Families |
|---|---|---|---|---|---|---|
| **EPT** | *Acrobeles*-like | 14 | 2, 3, 4, 5, 7, 8, 9, 11, 12, 13, 14, 15, 18, 22 | *Acrobeles* sp., *Acrobeloides* sp. | 1 | Cephalobidae |
| | *Acrobeloides* | 8 | 3, 4, 6, 8, 11, 14, 15, 17 | *Acrobeloides* sp. | 1 | Cephalobidae |
| | Alaimidae | 1 | 2 | *Alaimus* sp. | 1 | Alaimidae |
| | Aphelenchoidea | 4 | 1, 6, 7, 10 | Aphelenchoididae sp., *Aphelenchoides* sp. | 1 | Aphelenchoididae |
| | Dorylaimida | 9 | 2, 3, 4, 5, 6, 7, 12, 13, 14 | Dorylaimida sp., Dorylaimina sp., Discolaimium sp. | 4 | Aporcelaimidae, Pararhyssocolpidae, Qudsianematidae, Undefined |
| | *Panagrolaimus* | 3 | 5, 7, 19 | *Panagrolaimus* sp. | 1 | Panagrolaimidae |
| | **total** | **39** | | | **9** | |
| **ARO** | *Acrobeles*-like | 2 | 3, 8 | *Acrobeles* sp. | 1 | Cephalobidae |
| | *Acrobeloides* | 3 | 1, 19, 24 | *Acrobeloides* sp. | 1 | Cephalobidae |
| | Alaimidae | 1 | 2 | *Alaimus* sp. | 1 | Alaimidae |
| | Aphelenchoidea | 5 | 2, 13, 14, 15, 16 | *Aphelenchoides* sp. | 1 | Aphelenchoididae |
| | Dorylaimida | 2 | 11, 15 | Dorylaimida sp. | 1 | Aporcelaimidae |
| | *Panagrolaimus* | 3 | 5, 19, 20 | *Panagrolaimus* sp. | 1 | Panagrolaimidae |
| | Plectidae | 1 | 9 | *Plectus* sp. | 1 | Plectidae |
| | **total** | **17** | | | **7** | |
| **TDT** | *Acrobeles*-like | 5 | 1, 10, 17, 19, 21 | *Acrobeles* sp. | 1 | Cephalobidae |
| | *Acrobeloides* | 8 | 2, 6, 12, 13, 18, 19, 20, 21 | *Acrobeloides* sp. | 1 | Cephalobidae |
| | *Panagrolaimus* | 1 | 16 | *Panagrolaimus* sp. | 1 | Panagrolaimidae |
| | **total** | **14** | | | **3** | |
| **ALT** | *Acrobeles*-like | 1 | 16 | *Acrobeles* sp. | 1 | Cephalobidae |
| | *Acrobeloides* | 5 | 5, 10, 11, 16, 24 | *Acrobeloides* sp. | 1 | Cephalobidae |
| | Aphelenchoidea | 1 | 10 | *Aphelenchoides* sp. | 1 | Aphelenchoididae |
| | Dorylaimida | 1 | 10 | Dorylaimida sp. | 1 | Aporcelaimidae |
| | *Panagrolaimus* | 8 | 2, 5, 6, 11, 12, 13, 14, 15 | *Panagrolaimus* sp. | 1 | Panagrolaimidae |
| | Plectidae | 1 | 4 | *Plectus* sp. | 1 | Plectidae |
| | **total** | **17** | | | **6** | |
| **PAP** | *Acrobeloides* | 2 | 7, 24 | *Acrobeloides* sp. | 1 | Cephalobidae |
| | Aphelenchoidea | 1 | 5 | *Aphelenchoides* sp. | 1 | Aphelenchoididae |
| | Dorylaimida | 1 | 1 | Dorylaimida sp. | 1 | Aporcelaimidae |
| | *Panagrolaimus* | 5 | 1, 4, 16, 17, 18 | *Panagrolaimus* sp. | 1 | Panagrolaimidae |
| | **total** | **9** | | | **4** | |
| **LAG** | *Acrobeloides* | 1 | 22 | *Acrobeloides* sp. | 1 | Cephalobidae |
| | Plectidae | 1 | 9 | *Plectus* sp. | 1 | Plectidae |
| | **total** | **2** | | | **2** | |
| **SLH** | *Acrobeloides* | 1 | 19 | *Acrobeloides* sp. | 1 | Cephalobidae |
| | Alaimidae | 1 | 2 | *Alaimus* sp. | 1 | Alaimidae |
| | Dorylaimida | 2 | 9, 15 | Dorylaimida sp. | 2 | Aporcelaimidae, Dorylaimidae |
| | Plectidae | 6 | 1, 2, 4, 5, 6, 10 | *Plectus* sp. | 1 | Plectidae |
| | **total** | **10** | | | **5** | |
| **NCBI References** | *Acrobeles*-like | 4 | 3, 6, 17, 20 | | 2 | Cephalobidae, Drilonematidae |
| | *Acrobeloides* | 3 | 9, 23, 24 | | 1 | Cephalobidae |
| | Alaimidae | 1 | 1 | | 1 | Cephalobidae |
| | Aphelenchoidea | 6 | 3, 4, 8, 9, 11, 12 | | 1 | Aphelenchoididae, Aphelenchidae |
| | Dorylaimida | 4 | 1, 8, 15, 16 | | 3 | Qudsianematidae, Aporcelaimidae, Pararhyssocolpidae |
| | *Panagrolaimus* | 4 | 3, 8, 9, 10 | | 1 | Panagrolaimidae |
| | Plectidae | 5 | 3, 4, 7, 8, 9 | | 1 | Plectidae |

**Table S11. Colonizer persister category proportion of the families found in different sampling transects.**

| | Altiplano | Aroma | EaglePoint | Paposo | Salars | TotoralDunes |
|---|---|---|---|---|---|---|
| CP 1 | 14.3 | 14.3 | 12.5 | 11.1 | 9.1 | 28.6 |
| CP 2 | 42.9 | 42.9 | 37.5 | 33.3 | 27.3 | 28.6 |
| CP 3 | 0.0 | 0.0 | 0.0 | 0.0 | 18.2 | 0.0 |
| CP 4 | 28.6 | 28.6 | 37.5 | 33.3 | 36.4 | 28.6 |
| CP 5 | 14.3 | 14.3 | 12.5 | 22.2 | 9.1 | 14.3 |
| *CP1+CP2* | *57.2* | *57.2* | *50* | *44.4* | *36.4* | *57.2* |
| *CP4+CP5* | *42.9* | *42.9* | *50* | *55.5* | *45.5* | *42.9* |

**Table S12. Feeding types of the families found in the different sampling transects.**

| | Altiplano | Aroma | Eagle Point | Paposo | Salars | Totoral Dunes |
|---|---|---|---|---|---|---|
| Herbivores | 0.0 | 0.0 | 11.1 | 10.0 | 0.0 | 12.5 |
| Fungivores | 28.6 | 14.3 | 11.1 | 20.0 | 0.0 | 12.5 |
| Bacterivores | 42.9 | 57.1 | 44.4 | 20.0 | 45.5 | 37.5 |
| Predators | 14.3 | 14.3 | 11.1 | 20.0 | 27.3 | 25.0 |
| Unicelular eukaryotic feeders | 0.0 | 0.0 | 0.0 | 0.0 | 0.0 | 0.0 |
| Omnivores | 14.3 | 14.3 | 22.2 | 30.0 | 27.3 | 12.5 |

**Table S13. Geomorphological and climatic variables tested for the modeling approaches.**

| Variable | Description | Type | Kind | min | max |
|---|---|---|---|---|---|
| Response | Composition of genera | integer | Biotic | 0 | 6 |
| Predictive | Elevation | numeric | Geomorphological | 84 | 4341 |
| Predictive | Topographic complexity | numeric | Geomorphological | 1,52819 | 214,6917 |
| Predictive | Soil thickness | numeric | Geomorphological | 0 | 50 |
| Predictive | Curvature | numeric | Geomorphological | -3200 | 900 |
| Predictive | Rock type | integer | Geomorphological | 1 | 11 |
| Predictive | Latitude | numeric | Geomorphological | -27.79444 | -19.52667 |
| Predictive | Mean annual temperature | numeric | Climatic | 4.325012 | 21.75835 |
| Predictive | Range of temperature | numeric | Climatic | 2.700012 | 6.799988 |
| Predictive | Mean annual precipation | numeric | Climatic | 3.3 | 311.2 |
| Predictive | Range of precipitation | numeric | Climatic | 0.7 | 91.799995 |

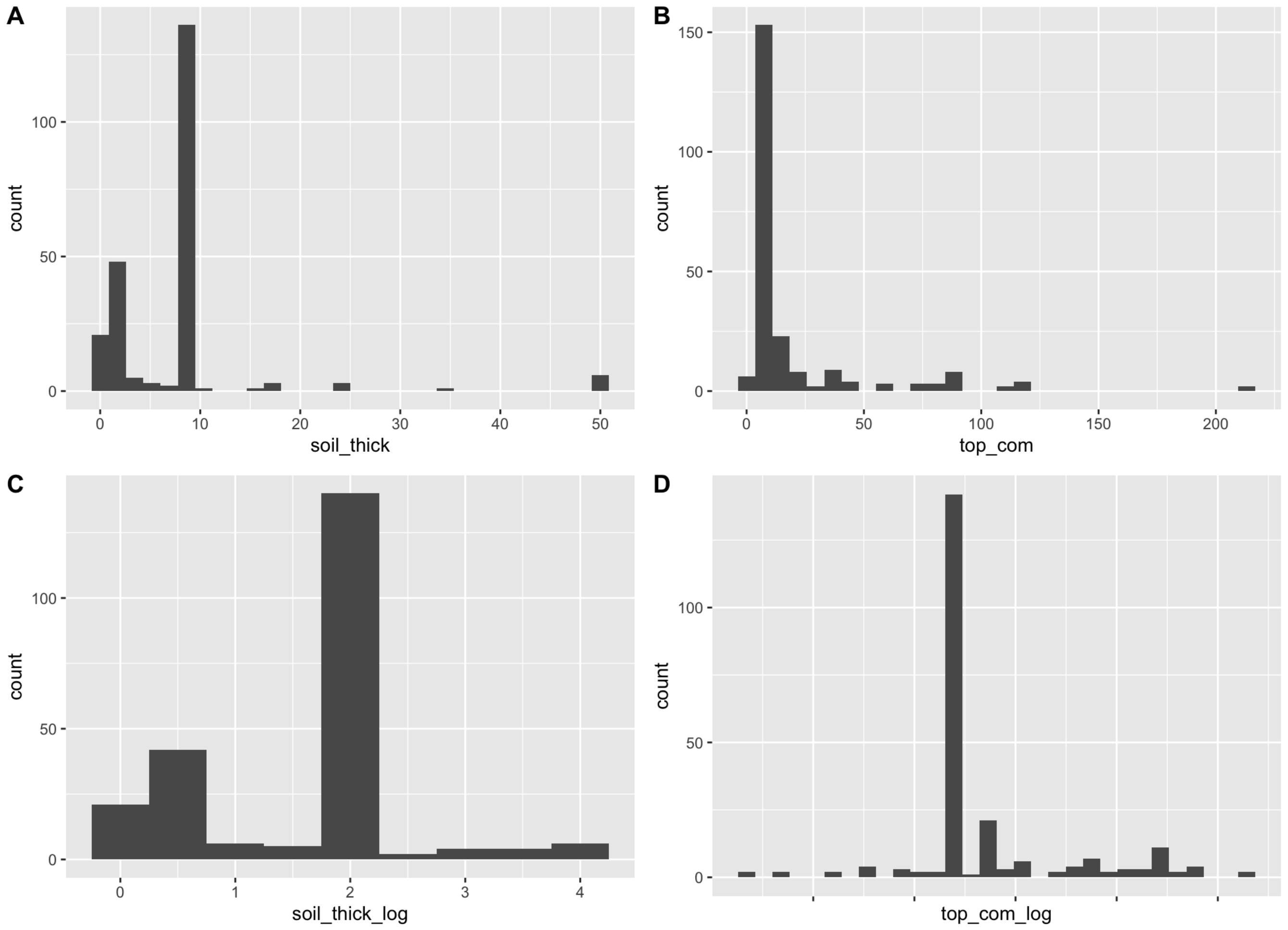


**Fig. S8.** Transformation of soil thickness (A and C) and topographic complexity (B and D) for implementation in generalised linear model.

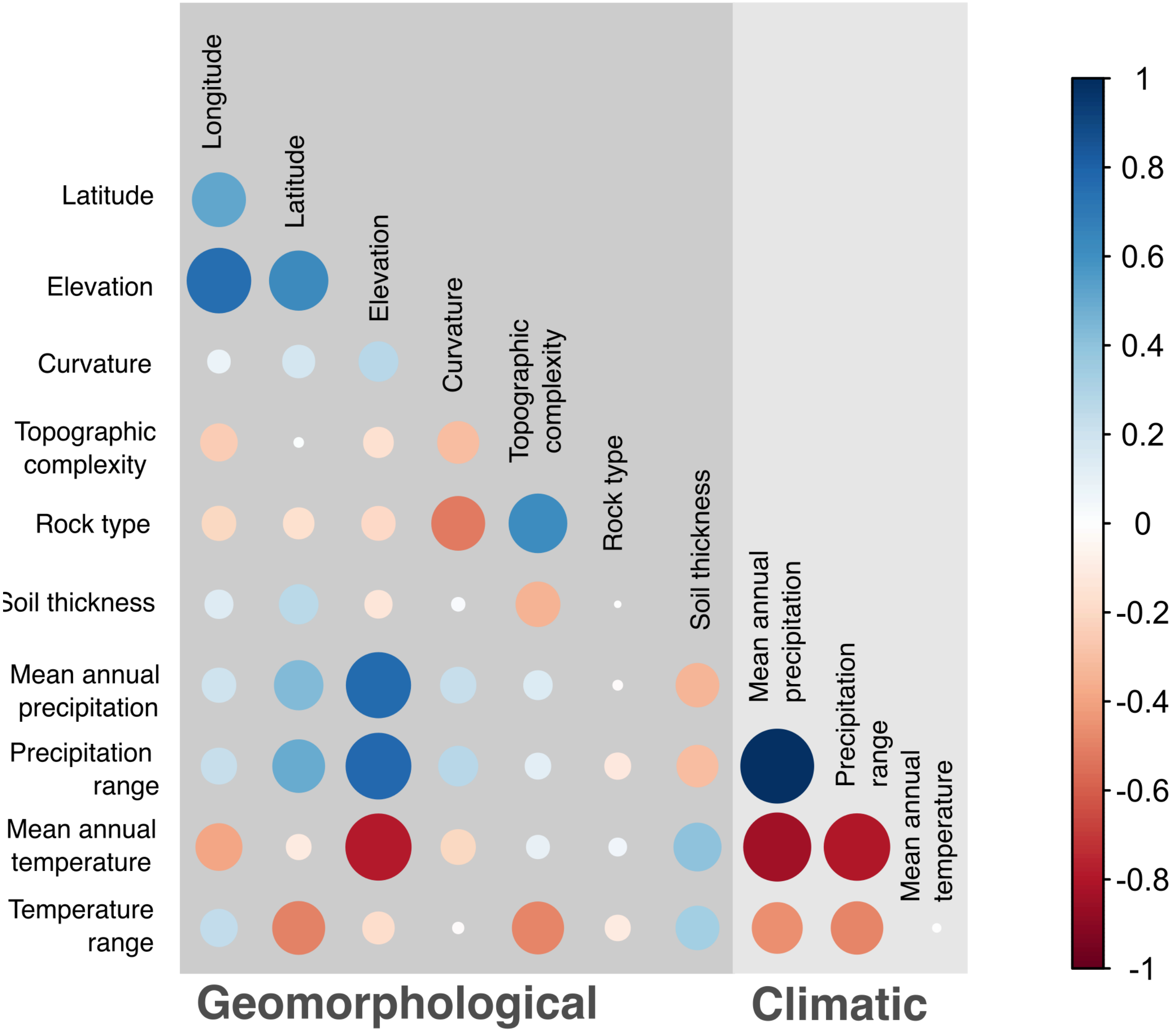


**Fig. S9.** Correlation of geographical and climatic variables considered as possible predictors in the Generalized Least Squares models(GLS), Linear mixed-effects models (LME) and Random forest approach to determine genera richness in the desert (Pearson method). Variables mean annual precipitation and precipitation range show a high positive correlation (0.9884, $p<0.05$). Similarly, mean annual temperature and mean annual precipitation exhibit a strong negative correlation (-0.8357, $p<0.05$). Additionally, mean annual temperature and precipitation range also display a high negative correlation (-0.7967, $p<0.05$). Based on these correlations, the variables selected for further analysis include latitude, elevation, curvature, topographic complexity, rock type, soil thickness, mean annual precipitation, and range of temperature.

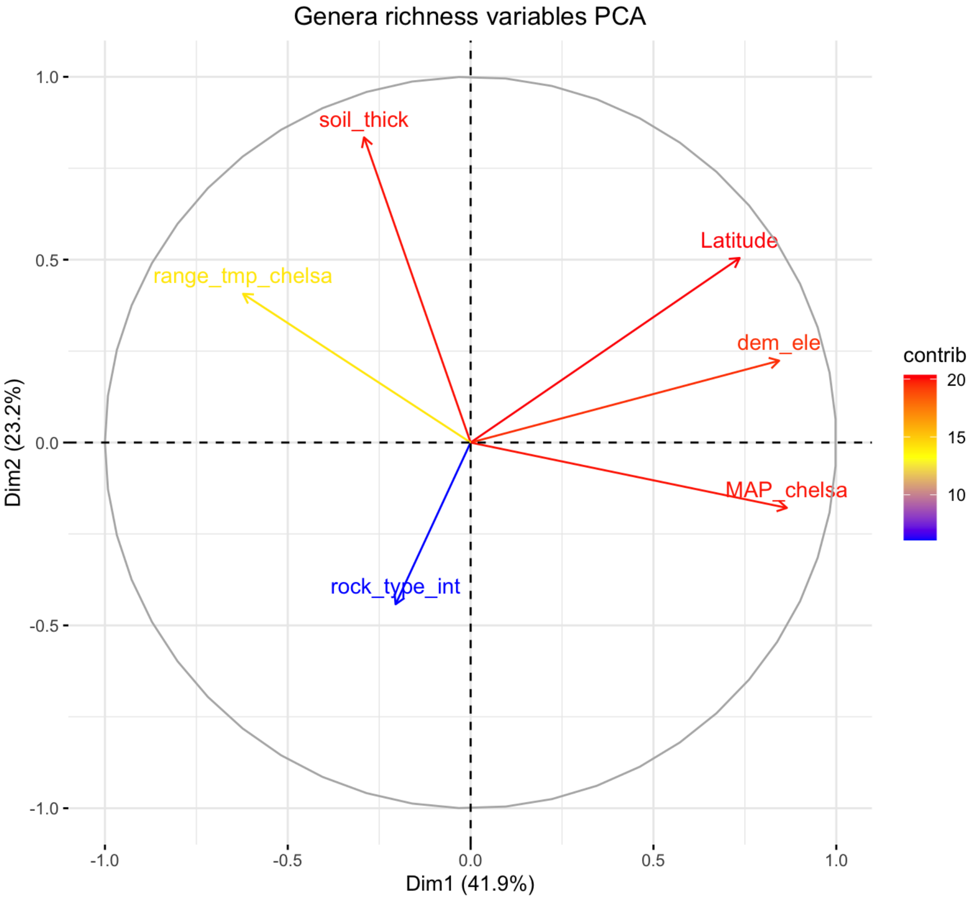


**Fig. S10.** Principal component analysis of variables to be used as predictor for genera richness in general linear models

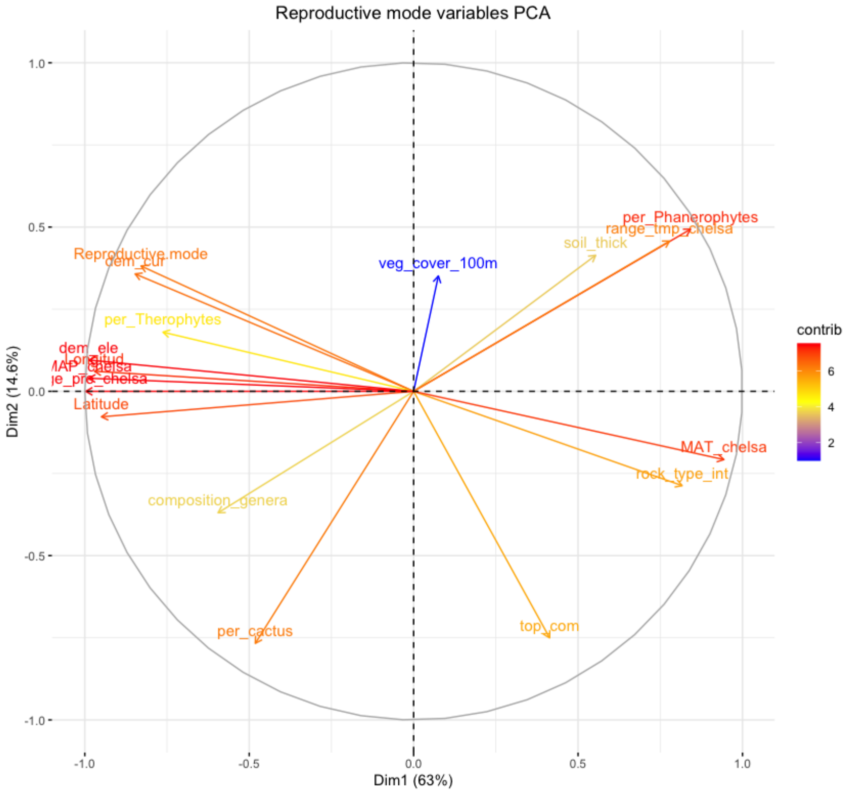


**Fig. S11.** Principal component analysis of variables to be used as predictor for reproductive mode in general linear models

**Table S14. Generalized least squares model specifications for genera richness displaying combination of variables tested. The best model (M_4_14), incorporation latitude and mean annual precipitation) is highlighted and was chosen according to the lowest BIC and highest akaike weight.**

| Model | Model specification using GLS | BICs | Akaike weights |
|---|---|---|---|
| M_1_0 | Latitude + dem_ele + rock_type_int + soil_thick_log + MAP_chelsa + range_tmp_chelsa | 439,3296 | 2,50E-03 |
| M_1_1 | Latitude + dem_ele + rock_type_int + soil_thick_log + MAP_chelsa | 435,7193 | 1,52E-02 |
| M_1_2 | Latitude + dem_ele + rock_type_int + soil_thick_log + range_tmp_chelsa | 426,1046 | 1,86E+00 |
| M_1_3 | Latitude + dem_ele + rock_type_int + MAP_chelsa + range_tmp_chelsa | 433,7931 | 3,98E-02 |
| M_1_4 | Latitude + dem_ele + soil_thick_log + MAP_chelsa + range_tmp_chelsa | 425,7154 | 2,26E+00 |
| M_1_5 | Latitude + rock_type_int + soil_thick_log + MAP_chelsa + range_tmp_chelsa | 422,8971 | 9,24E+00 |
| M_1_6 | dem_ele + rock_type_int + soil_thick_log + MAP_chelsa + range_tmp_chelsa | 436,6535 | 9,52E-03 |
| M_2_1 | Latitude + dem_ele + rock_type_int + soil_thick_log | 416,7643 | 1,98E+02 |
| M_2_10 | Latitude + soil_thick_log + MAP_chelsa + range_tmp_chelsa | 409,2474 | 8,51E+03 |
| M_2_11 | dem_ele + rock_type_int + soil_thick_log + MAP_chelsa | 452,9547 | 2,75E-06 |
| M_2_12 | dem_ele + rock_type_int + soil_thick_log + range_tmp_chelsa | 424,355 | 4,46E+00 |
| M_2_13 | dem_ele + rock_type_int + MAP_chelsa + range_tmp_chelsa | 437,3316 | 6,78E-03 |
| M_2_14 | dem_ele + soil_thick_log + MAP_chelsa + range_tmp_chelsa | 427,9819 | 7,27E-01 |
| M_2_15 | rock_type_int + soil_thick_log + MAP_chelsa + range_tmp_chelsa | 416,6406 | 2,11E+02 |
| M_2_2 | Latitude + dem_ele + rock_type_int + MAP_chelsa | 429,0705 | 4,22E-01 |
| M_2_3 | Latitude + dem_ele + rock_type_int + range_tmp_chelsa | 421,6383 | 1,73E+01 |
| M_2_4 | Latitude + dem_ele + soil_thick_log + MAP_chelsa | 426,5662 | 1,48E+00 |
| M_2_5 | Latitude + dem_ele + soil_thick_log + range_tmp_chelsa | 417,274 | 1,54E+02 |
| M_2_6 | Latitude + dem_ele + MAP_chelsa + range_tmp_chelsa | 424,88 | 3,43E+00 |
| M_2_7 | Latitude + rock_type_int + soil_thick_log + MAP_chelsa | 425,2025 | 2,92E+00 |
| M_2_8 | Latitude + rock_type_int + soil_thick_log + range_tmp_chelsa | 422,7985 | 9,71E+00 |
| M_2_9 | Latitude + rock_type_int + MAP_chelsa + range_tmp_chelsa | 413,251 | 1,15E+03 |
| M_3_1 | Latitude + dem_ele + rock_type_int | 414,6345 | 5,75E+02 |
| M_3_10 | Latitude + MAP_chelsa + range_tmp_chelsa | 403,9415 | 1,21E+05 |
| M_3_11 | dem_ele + rock_type_int + soil_thick_log | 438,3542 | 4,07E-03 |
| M_3_12 | dem_ele + rock_type_int + MAP_chelsa | 448,9546 | 2,03E-05 |
| M_3_13 | dem_ele + rock_type_int + range_tmp_chelsa | 432,5371 | 7,45E-02 |
| M_3_14 | dem_ele + soil_thick_log + MAP_chelsa | 444,0869 | 2,31E-04 |
| M_3_15 | dem_ele + soil_thick_log + range_tmp_chelsa | 411,4559 | 2,82E+03 |
| M_3_16 | dem_ele + MAP_chelsa + range_tmp_chelsa | 428,6293 | 5,26E-01 |
| M_3_17 | rock_type_int + soil_thick_log + MAP_chelsa | 430,1279 | 2,49E-01 |
| M_3_18 | rock_type_int + soil_thick_log + range_tmp_chelsa | 410,4978 | 4,55E+03 |
| M_3_19 | rock_type_int + MAP_chelsa + range_tmp_chelsa | 412,6911 | 1,52E+03 |
| M_3_2 | Latitude + dem_ele + soil_thick_log | 407,6717 | 1,87E+04 |
| M_3_20 | soil_thick_log + MAP_chelsa + range_tmp_chelsa | 403,3468 | 1,63E+05 |
| M_3_3 | Latitude + dem_ele + MAP_chelsa | 419,8891 | 4,16E+01 |
| M_3_4 | Latitude + dem_ele + range_tmp_chelsa | 412,7208 | 1,50E+03 |
| M_3_5 | Latitude + rock_type_int + soil_thick_log | 427,7891 | 8,01E-01 |
| M_3_6 | Latitude + rock_type_int + MAP_chelsa | 415,4604 | 3,81E+02 |
| M_3_7 | Latitude + rock_type_int + range_tmp_chelsa | 427,4303 | 9,58E-01 |
| M_3_8 | Latitude + soil_thick_log + MAP_chelsa | 411,7137 | 2,48E+03 |
| M_3_9 | Latitude + soil_thick_log + range_tmp_chelsa | 409,1147 | 9,09E+03 |
| M_4_1 | Latitude + dem_ele | 405,5521 | 5,40E+04 |
| M_4_10 | rock_type_int + soil_thick_log | 418,4105 | 8,71E+01 |
| M_4_11 | rock_type_int + MAP_chelsa | 424,454 | 4,24E+00 |
| M_4_12 | rock_type_int + range_tmp_chelsa | 422,2909 | 1,25E+01 |
| M_4_13 | soil_thick_log + MAP_chelsa | 424,3785 | 4,41E+00 |
| M_4_14 | soil_thick_log + range_tmp_chelsa | 401,2194 | 4,71E+05 |
| M_4_15 | MAP_chelsa + range_tmp_chelsa | 404,3134 | 1,00E+05 |
| M_4_2 | Latitude + rock_type_int | 423,1949 | 7,96E+00 |
| M_4_3 | Latitude + soil_thick_log | 417,6212 | 1,29E+02 |
| M_4_4 | Latitude + MAP_chelsa | 406,1876 | 3,93E+04 |
| M_4_5 | Latitude + range_tmp_chelsa | 419,5768 | 4,86E+01 |
| M_4_6 | dem_ele + rock_type_int | 434,9016 | 2,29E-02 |
| M_4_7 | dem_ele + soil_thick_log | 424,7641 | 3,63E+00 |
| M_4_8 | dem_ele + MAP_chelsa | 435,4706 | 1,72E-02 |
| M_4_9 | dem_ele + range_tmp_chelsa | 424,6992 | 3,75E+00 |

**Table S15. Linear mixed effect model specifications for genera richness displaying combination of variables tested using sampling locality as a random effect. The best model (M_4_15), incorporation latitude and mean annual precipitation) is highlighted and was chosen according to the lowest BIC and highest akaike weight.**

| Model | Model specification using LME, locality is used as a random effect | BICs | Akaike weights |
|---|---|---|---|
| M_1_0 | Latitude + dem_ele + rock_type_int + soil_thick_log + MAP_chelsa + range_tmp_chelsa | 422,6176095 | 4,74E-08 |
| M_1_1 | Latitude + dem_ele + rock_type_int + soil_thick_log + MAP_chelsa | 424,6243616 | 1,74E-08 |
| M_1_2 | Latitude + dem_ele + rock_type_int + soil_thick_log + range_tmp_chelsa | 419,0790102 | 2,78E-07 |
| M_1_3 | Latitude + dem_ele + rock_type_int + MAP_chelsa + range_tmp_chelsa | 417,7927052 | 5,29E-07 |
| M_1_4 | Latitude + dem_ele + soil_thick_log + MAP_chelsa + range_tmp_chelsa | 414,6804345 | 2,51E-06 |
| M_1_5 | Latitude + rock_type_int + soil_thick_log + MAP_chelsa + range_tmp_chelsa | 405,1242572 | 0,000297926 |
| M_1_6 | dem_ele + rock_type_int + soil_thick_log + MAP_chelsa + range_tmp_chelsa | 417,3134641 | 6,72E-07 |
| M_2_1 | Latitude + dem_ele + rock_type_int + soil_thick_log | 418,8969328 | 3,04E-07 |
| M_2_10 | Latitude + soil_thick_log + MAP_chelsa + range_tmp_chelsa | 397,1009353 | 0,016457014 |
| M_2_11 | dem_ele + rock_type_int + soil_thick_log + MAP_chelsa | 420,9095299 | 1,11E-07 |
| M_2_12 | dem_ele + rock_type_int + soil_thick_log + range_tmp_chelsa | 414,1289397 | 3,30E-06 |
| M_2_13 | dem_ele + rock_type_int + MAP_chelsa + range_tmp_chelsa | 413,378034 | 4,81E-06 |
| M_2_14 | dem_ele + soil_thick_log + MAP_chelsa + range_tmp_chelsa | 409,4868888 | 3,36E-05 |
| M_2_15 | rock_type_int + soil_thick_log + MAP_chelsa + range_tmp_chelsa | 400,7279595 | 0,002683816 |
| M_2_2 | Latitude + dem_ele + rock_type_int + MAP_chelsa | 418,6892497 | 3,38E-07 |
| M_2_3 | Latitude + dem_ele + rock_type_int + range_tmp_chelsa | 418,1167498 | 4,50E-07 |
| M_2_4 | Latitude + dem_ele + soil_thick_log + MAP_chelsa | 416,7165767 | 9,05E-07 |
| M_2_5 | Latitude + dem_ele + soil_thick_log + range_tmp_chelsa | 411,1751553 | 1,45E-05 |
| M_2_6 | Latitude + dem_ele + MAP_chelsa + range_tmp_chelsa | 409,8513978 | 2,80E-05 |
| M_2_7 | Latitude + rock_type_int + soil_thick_log + MAP_chelsa | 407,0560892 | 0,000113401 |
| M_2_8 | Latitude + rock_type_int + soil_thick_log + range_tmp_chelsa | 408,3053359 | 6,07E-05 |
| M_2_9 | Latitude + rock_type_int + MAP_chelsa + range_tmp_chelsa | 400,433734 | 0,00310916 |
| M_3_1 | Latitude + dem_ele + rock_type_int | 413,0202543 | 5,75E-06 |
| M_3_10 | Latitude + MAP_chelsa + range_tmp_chelsa | 392,3762298 | 0,174706001 |
| M_3_11 | dem_ele + rock_type_int + soil_thick_log | 418,2169625 | 4,28E-07 |
| M_3_12 | dem_ele + rock_type_int + MAP_chelsa | 414,7865542 | 2,38E-06 |
| M_3_13 | dem_ele + rock_type_int + range_tmp_chelsa | 417,9257209 | 4,95E-07 |
| M_3_14 | dem_ele + soil_thick_log + MAP_chelsa | 413,4520954 | 4,63E-06 |
| M_3_15 | dem_ele + soil_thick_log + range_tmp_chelsa | 406,3595877 | 0,000160642 |
| M_3_16 | dem_ele + MAP_chelsa + range_tmp_chelsa | 405,7096162 | 0,000222331 |
| M_3_17 | rock_type_int + soil_thick_log + MAP_chelsa | 406,4060312 | 0,000156955 |
| M_3_18 | rock_type_int + soil_thick_log + range_tmp_chelsa | 401,7714867 | 0,001592772 |
| M_3_19 | rock_type_int + MAP_chelsa + range_tmp_chelsa | 398,3985746 | 0,008601462 |
| M_3_2 | Latitude + dem_ele + soil_thick_log | 410,5359401 | 1,99E-05 |
| M_3_20 | soil_thick_log + MAP_chelsa + range_tmp_chelsa | 392,9100456 | 0,133780001 |
| M_3_3 | Latitude + dem_ele + MAP_chelsa | 410,6980986 | 1,84E-05 |
| M_3_4 | Latitude + dem_ele + range_tmp_chelsa | 410,1620367 | 2,40E-05 |
| M_3_5 | Latitude + rock_type_int + soil_thick_log | 409,3329669 | 3,63E-05 |
| M_3_6 | Latitude + rock_type_int + MAP_chelsa | 400,8435468 | 0,002533106 |
| M_3_7 | Latitude + rock_type_int + range_tmp_chelsa | 416,0325034 | 1,27E-06 |
| M_3_8 | Latitude + soil_thick_log + MAP_chelsa | 398,8873015 | 0,00673669 |
| M_3_9 | Latitude + soil_thick_log + range_tmp_chelsa | 400,5148918 | 0,002985519 |
| M_4_1 | Latitude + dem_ele | 404,61139 | 0,000385014 |
| M_4_10 | rock_type_int + soil_thick_log | 402,6054636 | 0,001049682 |
| M_4_11 | rock_type_int + MAP_chelsa | 400,2709829 | 0,003372749 |
| M_4_12 | rock_type_int + range_tmp_chelsa | 409,6545593 | 3,09E-05 |
| M_4_13 | soil_thick_log + MAP_chelsa | 398,7024322 | 0,007389081 |
| M_4_14 | soil_thick_log + range_tmp_chelsa | 393,8743207 | 0,082604081 |
| M_4_15 | MAP_chelsa + range_tmp_chelsa | 390,7481978 | 0,394303353 |
| M_4_2 | Latitude + rock_type_int | 410,2205922 | 2,33E-05 |
| M_4_3 | Latitude + soil_thick_log | 401,3565058 | 0,001960043 |
| M_4_4 | Latitude + MAP_chelsa | 392,625023 | 0,154270565 |
| M_4_5 | Latitude + range_tmp_chelsa | 408,2267192 | 6,32E-05 |
| M_4_6 | dem_ele + rock_type_int | 415,1069188 | 2,02E-06 |
| M_4_7 | dem_ele + soil_thick_log | 410,4611039 | 2,07E-05 |
| M_4_8 | dem_ele + MAP_chelsa | 407,2428007 | 0,000103293 |
| M_4_9 | dem_ele + range_tmp_chelsa | 410,4661639 | 2,06E-05 |

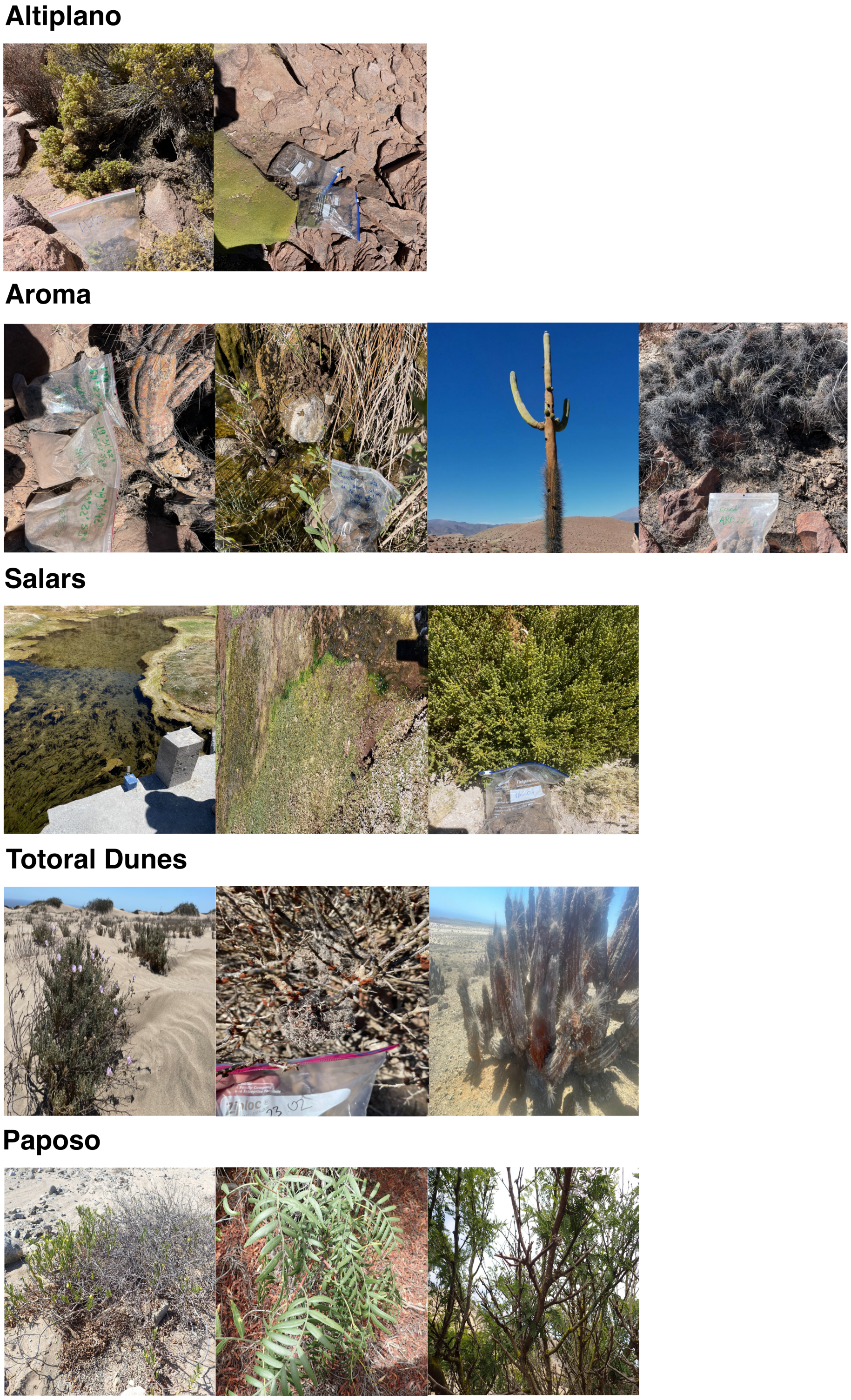


**Fig. S12.** Plants from the different sampling spots from which nematodes were cultured in the lab and which (nematodes') reproductive mode was assessed.

**Table S16. Variable importance for genera richness prediction in Random Forest. The most important variables given the highest %IncMSE and IncNodePurity are elevation (dem_ele), mean annual precipitation (MAP_chelsa) and latitude.**

| Variable | %IncMSE | IncNodePurity |
|---|---|---|
| Latitude | 14.51069 | 20.364106 |
| dem_ele | 17.62021 | 29.888430 |
| rock_type_int | 10.34478 | 9.472686 |
| soil_thick_log | 11.62650 | 13.886717 |
| MAP_chelsa | 17.81223 | 23.848930 |
| range_tmp_chelsa | 13.40530 | 17.503840 |

**Table S17. Variable importance for reproductive mode prediction in Random Forest. Elevation (dem_ele) is the most important variable according to the highest value of Mean Decrease accuracy and highest Mean Decrease Gini.**

| Variable | MeanDecreaseAccuracy | MeanDecreaseGini |
|---|---|---|
| dem_ele | 0.6176 | 1.6342 |
| MAP_chelsa | -1.2354 | 1.1678 |
| Latitude | -0.4347 | 1.2277 |

**Table S18. Lithological classes analyzed in the different modeling approaches from the global lithological map database GLiM.**

| Rock type integer | Lithological class | Implementation in code |
|---|---|---|
| 1 | Unconsolidated sediments (SU) | UncSed |
| 2 | Siliciclastic sedimentary rocks (SS) | SilRck |
| 3 | Pyroclastic rocks (PY) | PyrRck |
| 4 | Mixed sedimentary rocks (SC) | MxSRck |
| 5 | Carbonate sedimentary rocks (SM) | CarRck |
| 6 | Evaporites (EV) | EvaRck |
| 7 | Acid plutonic rocks (PA) | AciRck |
| 8 | Intermidiate volcanic rocks (VI) | InVRck |
| 9 | Basic volcanic rocks (VB) | BaVRck |
| 10 | Acidic plutonic rocks (PA) | AcPRck |
| 11 | Intermidiate plutonic rocks (PI) | InPRck |
| 12 | Basic plutonic rocks (PB) | BaPRck |
| 13 | Metamorphic rocks (MT) | MetRck |
| 14 | Water bodies (WB) | WatBod |
| 15 | Ice and Glaciers (IG) | IceGla |
| 16 | No data | nan |

**Table S19. Plant identification and estimation of vegetation coverage, percentage of phanerophytes, therophytes and cactus in the sampling areas from which isolated nematodes where cultured and which (nematodes') reproductive mode was assessed. The  represents a morphologically identified species without genetic data.**

| Sampling Location | Nematode genera | Nematode reproductive mode | Plant identification | Formation | Vegetation cover over 100m2 | % Phanerophytes (tree/shrub) | % Therophytes (herb) | % cactus |
|---|---|---|---|---|---|---|---|---|
| LAG.22.03 | Plectus | asexual | Fabiana ramulosa | Matorral bajo de altitud | 50 | 70 | 30 | 0 |
| LAG.22.01 | Acrobeloides | asexual | Fabiana ramulosa | Matorral bajo de altitud | 50 | 70 | 30 | 0 |
| SLH.23.01 | Plectus | asexual | Oxychloe andina | Salt flat | 90 | 0 | 100 | 0 |
| SLH.23.07 | Propanagrolaimus* | asexual | Oxychloe andina | Salt flat | 90 | 0 | 100 | 0 |
| ALT.22.08 | Panagrolaimus | asexual | Azorella compacta | Matorral bajo de altitud | 10 | 90 | 10 | 0 |
| ALT.22.04 | Panagrolaimus | asexual | Fabiana ramulosa | Matorral bajo de altitud | 30 | 70 | 30 | 0 |
| ARO.22.31 | Panagrolaimus | sexual | Browningia candelaris | Bosque espinoso | 10 | 30 | 60 | 10 |
| ARO.22.27 | Eucephalobus | asexual | Dead cactus | Bosque espinoso | 15 | 30 | 50 | 20 |
| ARO.22.19 | Plectus | asexual | Corryocactus brevistylus | Bosque espinoso | 5 | 0 | 0 | 100 |
| ARO.22.19 | Panagrolaimus | sexual | Corryocactus brevistylus | Bosque espinoso | 5 | 0 | 0 | 100 |
| ARO.22.05 | Acrobeloides | asexual | Tessaria absinthioides | Desierto absoluto | 50 | 95 | 5 | 0 |
| PAP.22.02 | Panagrolaimus | sexual | Eulychnia sp (maybe) | Matorral desertico | 40 | 50 | 5 | 45 |
| PAP.22.29 | Panagrolaimus | sexual | Solanum chilense | Matorral desertico | 15 | 80 | 20 | 0 |
| PAP.22.38 | Panagrolaimus | sexual | Prosopis chilensis | Matorral desertico | 80 | 80 | 20 | 0 |
| PAP.22.39 | Panagrolaimus | sexual | Schinos molle | Matorral desertico | 80 | 80 | 20 | 0 |
| PAP.22.17 | Acrobeloides | asexual | Skytanthus acutus | Matorral desertico | 10 | 80 | 20 | 0 |
| TDT.23.02 | Acrobeles | sexual | Dead shrub with lichens | Matorral desertico | 10 | 90 | 10 | |
| TDT.23.14 | Cervidellus | sexual | Dead sticks | Matorral desertico | 10 | 90 | 10 | 0 |
| TDT.23.16 | Acrobeles | asexual | Dead shrub with lichens | Matorral desertico | 15 | 90 | 10 | 0 |

**Table S20. Model specifications for reproductive mode using GLM. the preferred model is glm1 incorporating only elevation according to lowest BIC and highest akaike weight.**

| Model | Model specifications for reproductive mode using GLM | BICs | Wm |
|---|---|---|---|
| glM0 | dem_ele + MAP_chelsa + Latitude + range_tmp_chelsa | 29,340250 | 0,0408556 |
| glM1 | dem_ele | 24,488020 | 0,43632563 |
| glM2 | MAP_chelsa | 26,155350 | 0,1895635 |
| glM3 | Latitude | 26,303670 | 0,17601435 |
| glM4 | Latitude + dem_ele | 27,367550 | 0,10340207 |
| glM5 | range_tmp_chelsa | 30,089800 | 0,02731868 |

**Table S21. Model specifications for reproductive mode using GLMM acounting for locality as a random effect. the preferred model is glm1 incorporating only elevation according to lowest BIC and highest akaike weight.**

| Model | Model specifications for reproductive mode using GLMM, sampling locality is used as a random effect | BICs | Wm |
|---|---|---|---|
| glM0_glmer | dem_ele + MAP_chelsa + Latitude + range_tmp_chelsa | 32,23062 | 0,03960763 |
| glM1_glmer | dem_ele | 27,3784 | 0,44815349 |
| glM2_glmer | MAP_chelsa | 29,04573 | 0,19470216 |
| glM3_glmer | Latitude | 29,19404 | 0,18078573 |
| glM4_glmer | Latitude + dem_ele | 30,25792 | 0,10620508 |
| glM5_glmer | range_tmp_chelsa | 32,75021 | 0,03054591 |

**Supplementary analysis: correlation between geographical distance and mean annual precipitation.** In order to test if the result form the modeling approach yielding mean annual temperature as one of the most important drivers for genera richness was driven by geographical distances. To address this, we performed a mantel test between geographical distance of the localities and mean annual precipitation difference between localities (code implemented can be found on the CodeOcean capsule (10.24433/CO.6395549.v2) as Ecological estimates: Geodist_MAP. The result showed that there is no significant correlation between these two characteristics (*rho*= 0.1321, p= 0.24861, 719 permutations), discarding that geographical distance is correlated to precipitation, geographical distance does not reflect the precipitation patterns in our dataset.
